# Neural Correlates of Statistical Learning in Developmental Dyslexia: An Electroencephalography Study

**DOI:** 10.1101/2022.07.06.498909

**Authors:** Tatsuya Daikoku, Sebastian Jentschke, Vera Tsogli, Kirstin Bergström, Thomas Lachmann, Merav Ahissar, Stefan Koelsch

## Abstract

The human brain extracts statistical regularities from the surrounding environment in a process referred to as statistical learning. Recent behavioural evidence suggests that developmental dyslexia affects statistical learning. However, surprisingly few neurophysiological studies have assessed how developmental dyslexia affects the neural processing underlying statistical learning. In this study, we used electroencephalography to explore the neural correlates of an important aspect of statistical learning – sensitivity to transitional probabilities – in individuals with developmental dyslexia. Adults diagnosed with developmental dyslexia (n = 17) and controls (n = 19) were exposed to a continuous stream of sound triplets in which a few triplet endings were location deviants (i.e., were presented from an unexpected speaker direction) or statistical deviants (i.e., had a low transitional probability given the triplet’s first two sounds). Location deviants elicited a large location mismatch negativity (MMN), which was larger in controls than dyslexics. Statistical deviants elicited a small, yet significant statistical MMN (sMMN) in controls, whereas the dyslexic individuals did not exhibit a statistical MMN. These results suggest that the neural mechanisms underlying statistical learning are impaired in developmental dyslexia.

**Significance statement:** We assessed the neural correlates of statistical learning in individuals with developmental dyslexia. Statistical deviants, namely word endings with a low transitional probability (as compared to high probability transitions) elicited a small, yet significant statistical MMN in controls, whereas the dyslexic individuals did not exhibit a statistical MMN. Location deviants elicited a MMN, which was larger in controls than dyslexics. These results suggest that the neural mechanisms underlying statistical learning are impaired in developmental dyslexia.

## Introduction

The brain can identify statistical regularities in sequential information in a process known as statistical learning (Saffran, Aslin, & Newport, 1996) or implicit learning (Perruchet & Pacton, 2006). Statistical learning involves an implicit and innate mechanism by which the brain calculates the transitional probability of sequential information. This learning system is thought to play a crucial role in early language acquisition. For example, 8-month-old infants (Saffran, Aslin, & Newport, 1996) and neonates (Teinonen et al., 2009) can learn the probabilities of syllable transitions, thus enabling them to detect word boundaries and isolate single words in natural speech.

Developmental dyslexia impairs reading comprehension and spelling in language, and is thought to arise mainly from phonological perceptual difficulties (Snowling, 2000; Ramus et al., 2003; Vellutino et al., 2004). In addition, individuals with developmental dyslexia may further have a wide range of other nonlinguistic as well as linguistic impairments, including weakened auditory statistical learning abilities (Arciuli & Simpson, 2012; Du & Kelly, 2013; Evans, Saffran, & Robe-Torres, 2009; Howard, Howard, Japikse, & Eden, 2006; Menghini, Hagberg, Caltagirone, Petrosini, & Vicari, 2006; Vicari et al., 2005) and non-linguistic perceptual processing (Ahissar, Protopapas, Reid, & Merzenich, 2000; Christmann, Lachmann, & Steinbrink, 2015; Giraud & Ramus, 2013; McAnally & Stein, 1996; Sperling, Lu, Manis, & Seidenberg, 2005). Since phonological deficits can also be observed in people without developmental dyslexia (Huettig et al., 2017), and not all children with developmental dyslexia show phonological processing deficits (Lachmann & van Leeuwen, 2008; Morris et al., 1998), phonological deficits may not automatically lead to developmental dyslexia. Thus, at least in some individuals, developmental dyslexia may also have other causes.

One possible cause is a more domain-general statistical learning deficit, not specific to phonological processing. Particularly, a great deal of evidence has shown that auditory statistical learning is impaired in dyslexic children, adolescents, and adults (Gabay et al., 2015; Dobo et al., 2021; Kahta et al., 2019; Vandermosten et al., 2019), although some other studies suggest that children with and without developmental dyslexia had no difference in statistical learning performance (Witteloostuijn et al., 2019). Despite much behavioral evidence on statistical learning deficits, the underlying neural mechanisms of these deficits remain unclear. This study aimed to investigate the neural correlates of statistical learning in individuals with developmental dyslexia using electroencephalography (EEG), which exhibits a high sensitivity to capture statistical learning even when behavioural measures may not indicate learning effects (Koelsch, Busch, Jentschke, & Rohrmeier, 2016).

EEG can be used to measure neural activity during statistical learning. Particularly, statistical learning is reflected in the electric brain responses to stimulation (event-related potentials, ERPs). When the brain encodes the statistics of stimulus sequences (e.g. regularities occurring within stimulus sequences), it predicts high-probability stimuli, which is associated with an inhibited ERP response to the predicted stimuli (compared with unpredicted, i.e., irregular or less regular, stimuli). Statistical learning effects can thus manifest as differences in ERP response amplitudes for expected and unexpected stimuli (for review, see Daikoku, 2018). Many studies have suggested that statistical learning is reflected in both early ERP components such as auditory brainstem response (ABR; Skoe, et al., 2015), P50 (Daikoku et al., 2017; Paraskevopoulos et al., 2012), N100 (Sanders et al., 2002), and mismatch negativity (MMN; Koelsch et al., 2016; Moldwin, Schwartz, & Sussman, 2017) as well as later ERP components such as the P200 (Balaguer et al., 2007; Cunillera et al., 2006) and theN400: (François, et al., 2013). Typically, a mismatch negativity (MMN) is elicited in response to physical deviants in oddball sequences. In such experiments, a series of standard stimuli is interspersed with physical deviants (oddballs; e..g, sound differing in pitch or location deviants; Christmann, Lachmann, & Berti, 2014; Garrido et al., 2008; Rinne, Antila, & Winkler, 2001; Sussman, Winkler, & Schröger, 2003; Winkler & Czigler, 2012). For example, if several sounds are presented from the right side, a sound presented on the left side elicits an MMN, which is generated mainly in the auditory cortex (Garrido et al., 2008).

Because developmental dyslexia is related to sensory processing dysfunctions, including those of the auditory cortex (Clark et al., 2014; Goswami, 2014; for a review see Gu & Bi, 2020), the MMN has been used to investigate the neural basis of developmental dyslexia (Kujala et al., 2000). A reduced MMN amplitude in children (Lachmann et al., 2005; for an overview see Bishop, 2007) and young adults (Schulte-Körne et al., 2001) with developmental dyslexia reflected impaired performance in syllable and tone discrimination and impaired tuning to native language speech representations (Bruder et al., 2011). The MMN has therefore been suggested as a neurophysiological endophenotype for developmental dyslexia (Neuhoff et al., 2012). However, there is also one study suggesting that only certain aspects of auditory processing may be affected: whereas the pitch MMN was shown to be impaired in that study, was the location MMN enhanced (Kujala et al., 2006). Notably, such MMN effects to physical changes do not require statistical learning because the perceptual regularities underlying the generation of the classical MMN can be extracted on a moment-to-moment basis, such as a series of stimuli coming from the right side interrupted by a stimulus from the left side. However, statistical learning can also be reflected in the MMN (François, Cunillera, Garcia, Laine, & Rodriguez-Fornells, 2017; Koelsch et al., 2016; Moldwin et al., 2017). Tsogli, Jentschke, Daikoku, and Koelsch (2019) presented sequences of tone triplets that could contain location deviants (comprising stimuli from an irregular location) and statistical deviants (comprising triplet endings with a low probability given the two preceding triplet items; this low transition probability could only be represented based on statistical learning, i.e., after extensive exposure to many triplets). Both statistical and location deviants elicited prominent mismatch ERP responses approximately 150–250 ms after stimulus onset. ERP effects elicited by statistical deviants are termed statistical MMN (sMMN; Koelsch et al., 2016) to distinguish them from the MMN to physical deviance, e.g., elicited by location deviants (Paavilainen, Karlsson, Reinikainen, & Näätänen, 1989; Sams, Paavilainen, Alho, & Näätänen, 1985). In contrast to the MMN eliecited by physical deviance, the elicitation of the sMMN does not occur on a moment-to-moment basis. Instead, it requires a more extended learning period to encode the underlying statistical regularities and to store them in long-term memory. Thus, the sMMN is suitable to investigate acquisition of knowledge regarding the statistical regularity of sound sequences.

This study investigated how developmental dyslexia affects the classical MMN in response to location deviants and the sMMN in response to statistical deviants in adults. Assuming that dyslexia adversely affects the inferences of sensory statistics in the auditory cortex (Jaffe-Dax, Kimel, & Ahissar, 2018; Lieder et al., 2019), we hypothesised that statistical and location deviants would elicit weaker MMNs in participants with developmental dyslexia than the controls. Confirmation of this hypothesis would provide evidence that developmental dyslexia is associated with both auditory processing dysfunction and statistical learning of auditory sequences.

## Materials and Methods

### Participants

Twenty-one adults diagnosed with developmental dyslexia (11 females, mean age = 26 years, SD = 5.3) and 20 age- and gender-matched control participants without a diagnosis of dyslexia (14 females, mean age = 26 years, SD = 3.1) were screened for eligibility to participate in this study. We excluded one individual from the developmental dyslexia group and one from the control group because both performed in the nonverbal intelligence test Standard Progressive Matrices (Raven & Court, 1998) with an IQ score < 70 (two SD below the normal range). Three other individuals with developmental dyslexia were excluded because they did not receive a clear diagnosis in childhood. After these exclusions, our study sample included 17 adults with developmental dyslexia and 19 control participants (Table 1), who all met the following inclusion criteria: German as the native language, right-handedness (Edinburgh Inventory; Oldfield, 1971), no history of neurological or audiological disorders, no diagnosis of a general or specific language impairment, no mental retardation, and no formal musical training for more than 5 years (beyond regular school lessons).

**Table 1.**
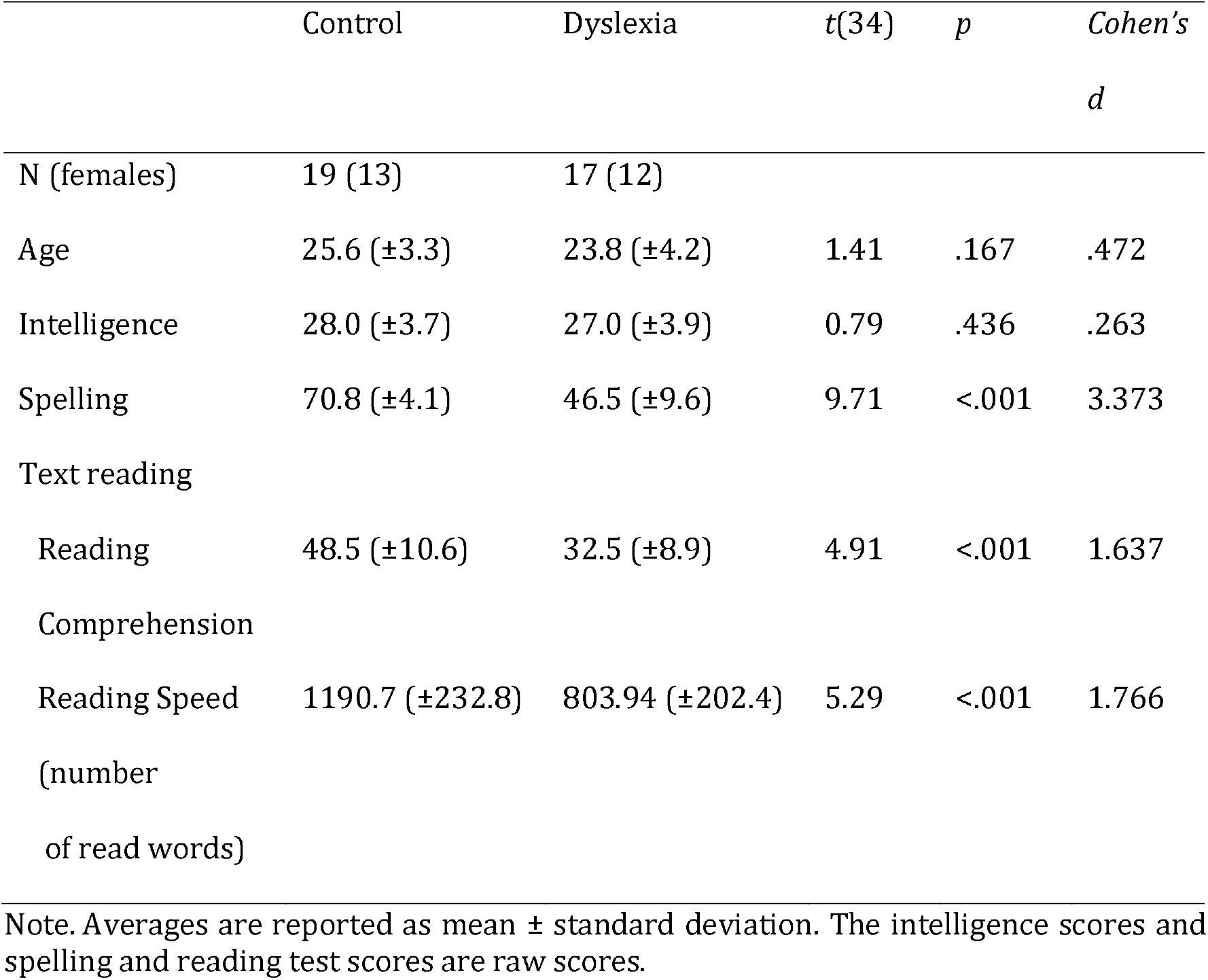
Demographic and diagnostic data (raw scores) in the control and dyslexia groups.

The study protocol was approved by the Ethics Committee of the Max-Planck-Institute (approval number: 2018/352). All participants were informed of the purpose of the study and the procedures in place to ensure their safety and the confidentiality of their personal data. They all provided written informed consent to participate in this study.

### Evaluation of Spelling and Reading Skills

Spelling skills were assessed with the Rechtschreibtest (Ibrahimović & Bulheller, 2013). In this test, participants were asked to fill in the missing words of a text, mainly composed of irregular German words, that a skilled and German native speaking experimenter read aloud. Silent text reading speeds and reading comprehension skills were assessed with the Lesegeschwindigkeits-und Verständnistest für die Klassen 5–12 (LVGT 5–12; Schneider, Schlagmüller, & Ennemoser, 2017). In this test, participants were asked to read as much of the text as possible within 4 min and to fill in each gap in the text with one of the three possible options.

### Stimuli

#### Sounds

*We* used the same stimuli and sequences as in a previous study (Tsogli et al., 2019). Each sound consisted of a Shepard tone (Shepard, 1964), combined with the sound produced by one of six different percussion instruments (i.e., a surdo, a tambourine, agogô bells, a hi-hat, castanets, or a woodblock). We obtained the percussive sounds from the Philharmonia Orchestra website (http://www.philharmonia.co.uk/explore/sound_samples). We used six distinct

Shepard tones based on six frequencies (i.e., F_3_ [174.61 Hz], G_3_ [196.00 Hz], A_3_ [220.00 Hz], B_3_ [246.94 Hz], C□_4_ [277.18 Hz], and D□_4_ [311.13 Hz]), each tone resulting from the superposition of nine sinusoidal components spaced an octave apart. The specific combinations of Shepard tones and percussive sounds were counterbalanced across participants. Examples of sounds are provided in Appendix A.

Another set of six sound combinations was created for a practice phase at the start of each experiment. These sounds were similar to those used in the main experiment but differed in terms of the frequencies providing bases for Shepard tones (i.e., E_3_ [164.81 Hz], F□_3_ [184.99 Hz], G□_3_ [207.65 Hz], A□_3_ [233.08 Hz], C_4_ [261.62 Hz], and D_4_ [293.66 Hz]) and the percussive sounds used (i.e., the sounds of a woodblock, a tambourine, agogô bells, castanets, a hi-hat, and a bass drum). An additional target Shepard tone based on C□_5_ (554.37 Hz), that did not have an accompanying percussive sound, was used for a cover task and a passive listening component of the experimental procedure (see Experimental Procedure).

All sound stimuli had a tone duration of 220 ms, including rising and falling periods of 10 ms and 20 ms, respectively, a constant loudness, and a sampling frequency of 44,100 Hz with 16 bit resolution.

#### Triplet Sequences

The stimuli descibed above, hereafter referred to as sounds A to F, were combined into sound triplets. Each 220 ms sound was followed by an 80 ms pause, such that the total duration of each triplet was 900 ms. As shown in Figure 1, sounds A and B and sounds C and D were paired to create two distinct two-sound sequences (i.e., AB and CD) that served as the first two sounds of each triplet, which are hereafter referred to as the triplet roots. Sounds E and F were the sounds that could be used as the last sound of a triplet, which is hereafter referred to as the triplet ending. Combining the two roots and two triplet endings yielded four possible triplets (i.e., ABE, ABF, CDE, and CDF). The assignment of different sounds to roles as triplet roots or endings was counterbalanced across participants as a way of ensuring that any possible acoustical differences between sounds would be cancelled out through particpants and not bias the neural responses of interest

**Figure 1.**
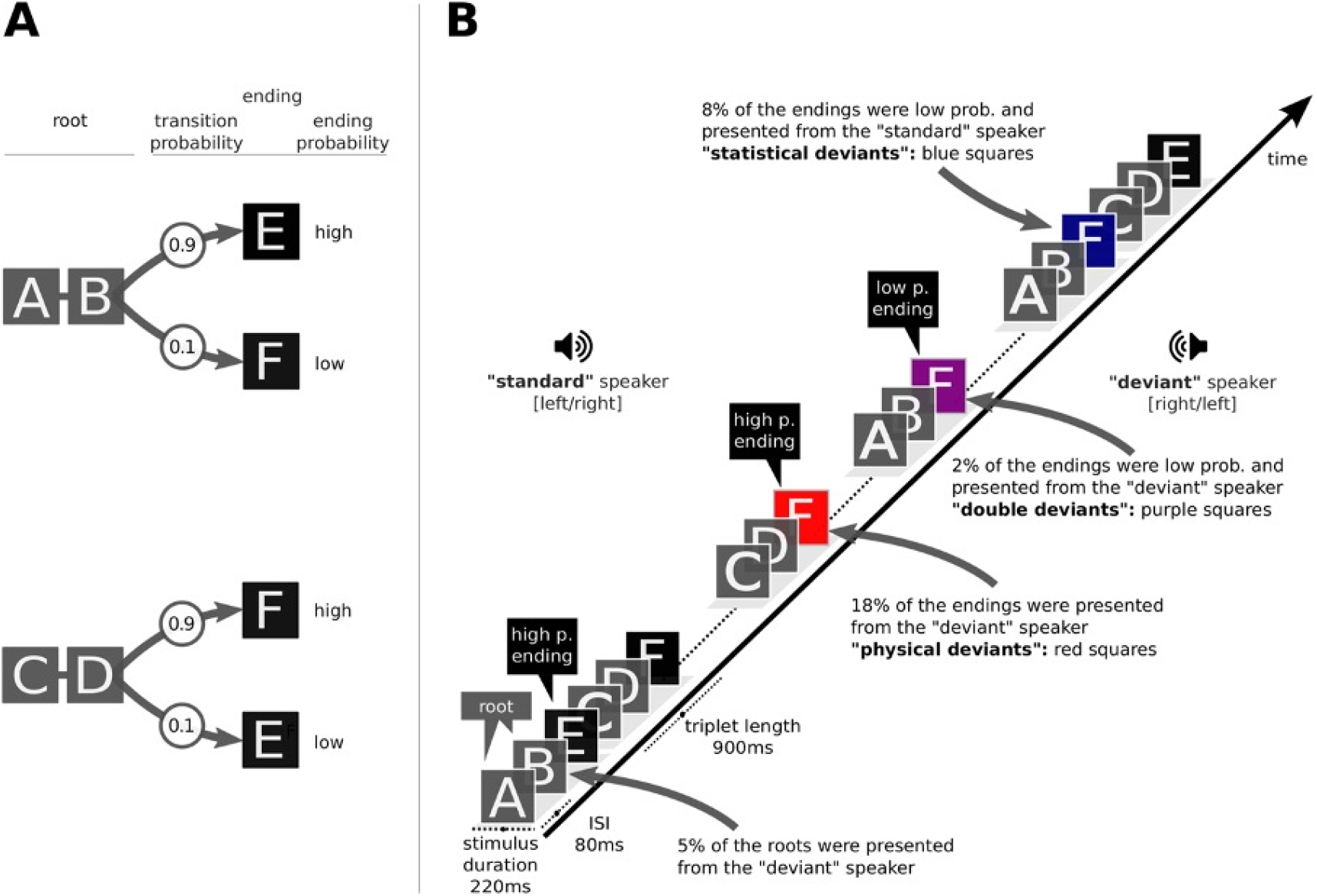
Triplets and stream in this study. **(A)** Four types of triplets generated from six different sounds (designated with letters from A to F) produced by pairing a Shepard tone with various percussive sounds. Each triplet consists of a root containing two conserved sounds (AB or CD) and a triplet ending with a high (90%) or a low (10%) transitional probability given the triplet’s root. Each triplet root has a 50% probability of occurring regardless of the ending sound of the previous triplet. (B) Example of a possible triplet sequence including standard triplets (triplet endings in black boxes), triplets with a statistically deviant ending (triplet endings in blue boxes) or a location deviant ending (triplet ending in red box), and a triplet with a doubly deviant ending (i.e., statistically deviant and location deviant; triplet ending in purple box). Reprinted, with permission, from Tsogli et al. (2019). Abbreviations: ISI, inter-stimulus interval; p. & prob., probability.

##### Exposition Sequences

The sequence of sounds used during an exposition phase comprised 400 sound triplets with a total sequence duration of about 6 min. The triplets were presented in a pseudo-randomised order with no two sequentially adjacent triplets being identical. Each of the two roots had an equal probability of occurring in a given triplet regardless of the ending sound of the previous triplet. Each statistically deviant triplet was followed by at least 3 triplets that were not statistically deviant.

Each sound stimulus was presented from either a speaker to the participant’s right or a speaker to the participant’s left. These speakers were positioned at 60° angles in the azimuthal plane. For each participant, one side was pseudo-randomly selected as the standard side for stimuli to be presented from, and the other side was the deviant one. The lateralization of the stimuli was balanced across blocks and counterbalanced between participants, and whether the location was “standard” or “deviant” was considered in the data analyses. For the triplet root sounds (i.e., sounds A, B, C, and D), 95% of the stimuli were presented from the standard side and the remaining 5% from the deviant side. For the triplet endings (i.e., sounds E and F), 80% of the stimuli were presented from the standard side and the remaining 20% from the deviant side. The triplets with endings presented from the deviant side were considered location deviants.

To generate statistical deviants, we set distinct probabilities for a transition (i.e., transitional probability) from a given root to a given ending within a triplet. The sensitivity to transitional probability is one of the important aspects of statistical learning mechanisms (Perruchet & Pacton, 2006; Saffran, Aslin, & Newport, 1996). The transitional probability for a given triplet ending was either 90% or 10% depending on root identity (Figure 1a). Triplets containing low-probability root-to-ending transitions were considered statistical deviants. Each statistically deviant triplet was followed by ≥3 triplets that were not statistically deviant. The location and statistical deviance created four triplet categories (Table 2). That is, standard triplets, which accounted for 72% of all triplets, were neither location deviant nor statistically deviant. The remaining 28% of triplets could be statistically deviant only (8% of all triplets), location deviant only (18% of all triplets), or both statistically deviant and location deviant (2% of all triplets).

**Table 2.**
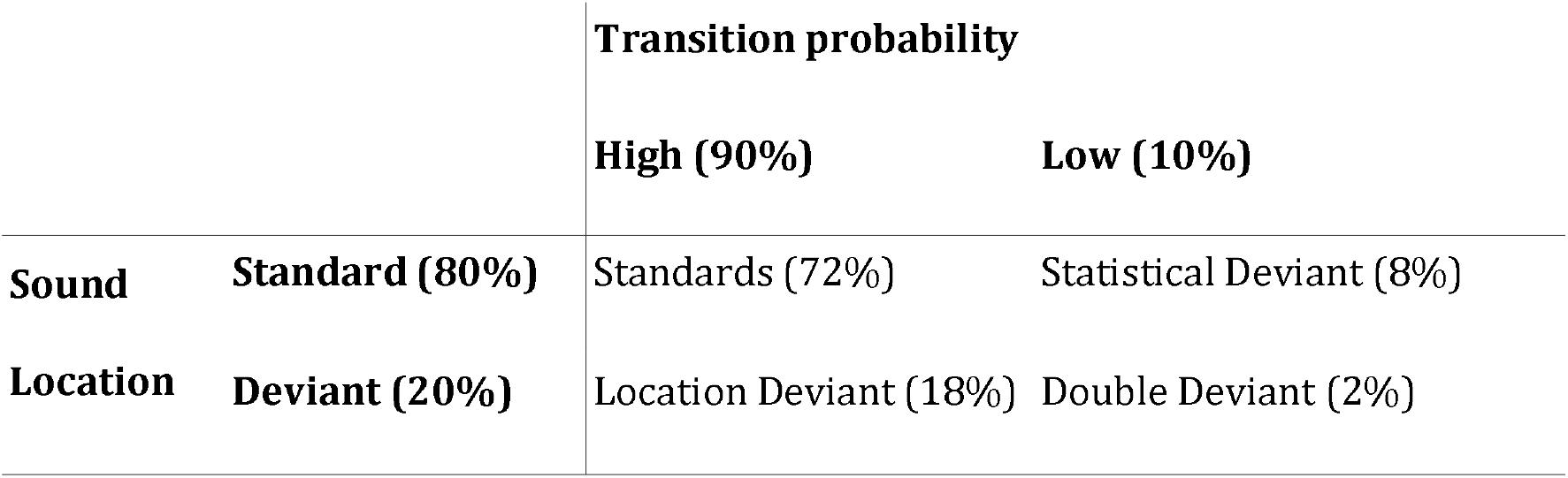
The 2×2 types of triplet endings based on location and statistical constraints.

##### Behavioural Testing Sequences

Each behavioural testing phase of the experiment consisted of twelve trials, with each trial involving a pair of triplets separated by a 335-ms pause. The triplets in each pair had the same root but different endings with low vs. high transition probabilities, respectively (i.e., statistically deviant vs. not deviant). The order in which the statistically deviant and standard triplets were played was counterbalanced across trials. Varying the order in which the same-root triplets within a pair were played created four sequentially distinct pairings (i.e., ABE to ABF, ABF to ABE, CDE to CDF, and CDF to CDE). Each sequentially distinct pairing was played three times during each behavioural testing phase, and consecutively played pairings had alternating triplet roots (i.e., each AB pairing was followed by a CD pairing that was in turn followed by an AB pairing). During behavioural testing phases, all triplets were played from both speakers, i.e., the task focussed on statistical regularity whereas location was not a factor.

### Experimental Design

The participants completed a multi-stage experiment with six blocks while undergoing EEG monitoring (see ‘Collection and Analysis of EEG Data’) inside an electromagnetically shielded chamber. Immediately before the experiment, the participants were provided with instructions concerning the procedures for the experiment’s different phases (see below). To ensure that only implicit learning could occur, the instructions did not include any description of the possible location or statistical deviance of the triplets. The participants then completed a 1-min practice session in which they were asked to press a key as soon as possible after hearing a target sound (i.e., C□_5_ [554.37 Hz]). If necessary, each participant repeated the practice session until they had correctly pressed the key after 80% of the target sound presentations.

Each of the six blocks included an exposition phase comprising the passive listening of the sequence of 400 triplets and a subsequent behavioural testing phase in which the participants performed actions based on a triplet sequence. During the exposition phase, participants were instructed to react to the high-pitched tones (cover tasks, 0.67% of all tones) while they were exposed to the sequence of 400 triplets described above (see Exposition Sequences). At the same time, they were watching a silent movie (nature or wildlife documentaries) played on a monitor in front of them. While the cover task was not particularly demanding (e.g., in terms of attentional resources) it minimized the possibility of the participants intentionally focusing their attention on statistical properties.

During a behavioural testing phase, the participants listened to paired statistically deviant and standard triplets (see Behavioural Testing Sequences). After each pair of triplets, a participant was asked to choose which triplet in the pair sounded more familiar and to rate their confidence in a given answer on a scale ranging from 1 (no certainty) to 5 (certainty).

### Acquisition and Analysis of EEG Data

*We* obtained 64-channel EEG data (Brain Amp, Brain Products, Munich, Germany) through cap-mounted electrodes placed over the participants’ scalps in accordance with the extended international 10-20 system. The left mastoid electrode was used as reference electrode and the neck electrode was used as ground electrode. The electrodes were clustered into six regions of interest: a frontal left region (F7, F5, F3, FT7, FC5, and FC3), a frontal middle region (F1, FZ, F2, FC1, FCZ, and FC2), a frontal right region (F8, F6, F4, FT8, FC6, and FC4), a central left region (T7, C5, C3, TP7, CP5, and CP3), a central middle region (C1, Cz, C2, and CPZ), and a central right region (T8, C6, C4, TP8, CP6, and CP4). Horizontal and vertical electro-oculograms were recorded bipolarly through electrodes placed at the outer canthi of the eyes and above and below the right eye. Electrode impedance was kept < 5 kΩ. Signals were recorded with a 0.25–1,000-Hz bandpass filter and a 500-Hz sampling rate.

EEG data were analysed in EEGLAB 13 (Delorme & Makeig, 2004) in MATLAB R2Ol8b (The MathWorks, Natick, Massachusetts). Continuous raw data files were re-referenced to the algebraic mean of the left and right mastoid electrodes and filtered with a 0.5-Hz high-pass filter and a 30-Hz low-pass filter implemented with finite impulse response designs and Blackman windows of 550 points and 2,750 points, respectively. Channels with excessive noise were identified through visual inspection and interpolated when necessary. The mean number of interpolated channels per participant was 0.22. Independent component analysis was used for linear decomposition of continuous data to remove the contributions of artefacts affecting scalp sensors (e.g. slow drifts, eye blinks or movement, and muscle artefacts). Epochs were removed from further analyses if the amplitude changes exceeded ±45 μV in any channels, including the electro-oculograms (less than 10% of the trials). The epochs of target stimuli (cover tasks) were removed in the analysis. In the end, 97.2% (SD±2.6%) and 94.8% (SD±8.0%) was preserved in the dyslexic and control groups, respectively. We also performed student’s t-test between groups. There were no significant differences (t(37), 0.69, p=0.50). Selective response averaging was conducted separately for standard triplets, triplets with a location deviant (but not with a statistical deviant), triplets with a statistical deviant (but not a location deviant), and triplets with both a location deviant and a statistical deviant (see ‘Exposition Sequences’).

Averages were computed using a 100 ms pre-stimulus baseline. We directly addressed the hypothesis of the present study, focusing on the average MMN amplitudes measured within a 150–250 ms time window. In addition, we analysed an obvious positive component approximately 100–140 ms after the onset of the stimuli, henceforth referred to as P120. That is, we also investigated the average amplitudes within a 100–140 ms time window.

### Statistical Analysis

#### Between-Group Comparisons of Participant Characteristics

Statistical analyses were conducted using jamovi version 1.2 (The jamovi Project, 2020). We used Bonferroni-corrected t-tests (dividing 0.05 by the number of tests) when comparing demographic characteristics and scores on tests of intelligence, spelling skills, and reading abilities between the dyslexia and the control group.

#### Analyses of Exposition Phase Data

*We* used separate ANOVA models to analyse the effects of stimulus deviance on P120 and MMN effect amplitudes during the exposition phases. The sMMN response is had a smaller amplitude size compared with the location MMN. Further, the sMMN is a relatively new ERP component, for which the neural mechanism has not yet been elucidated in detail. To determine those neural generators, studies using fMRI or MEG would be most suited, however, such studies still are lacking despite many studies using EEG methodology. Thus, this study directly compared between standard and deviant but did not use the subtraction waveform that is typically used in MMN studies. One ANOVA model featured the within-participant factor of sound location (i.e., location standard triplet endings or location deviant triplet endings), and the other featured the within-participant factor of transitional probability (i.e., statistically standard triplet endings or statistically deviant triplet endings). Both ANOVA models included the between-participants factor of groups (i.e., the developmental dyslexia vs. the control group) and three within-participant factors: the distinction between ERP responses in the frontal vs. central areas of the brain; the distinction between ERP responses in the left, medial, and right areas of the brain (i.e., ERP response lateralisation); and the different experimental blocks. The posterior region was not included because of low amplitude sizes. To boost our signal-to-noise ratio, our ANOVA models included three blocks rather than the actual six by merging the data from pairs of blocks.

We selected *p* < 0.05 as our threshold for statistical significance and used a false discovery rate method for the post-hoc testing of significant effects. To determine whether the Rechtschreibtest and LVGT 5–12 scores correlated with the P120 and MMN effect amplitudes, we calculated Pearson correlation coefficients.

#### Analyses of Behavioural Testing Phase Data

*We* used a two-tailed t-test to determine whether the frequency of correct answers during behavioural testing exceeded chance levels (i.e., >.5). We also used ANOVA models to compare the dyslexia and control groups in terms of response accuracies and reaction times (RŢ), and included experimental blocks as a within-subject factor in these analyses. As in our analyses of exposition phase data, we assumed three blocks rather than the actual six in order to boost our signal-to-noise ratios. We selected *p* <.05 as our threshold for statistical significance and used Bonferroni correction for the post-hoc testing of significant effects. We conducted Pearson correlation analysis to determine whether response accuracy rates correlated with confidence ratings and dyslexia test scores.

## Results

### Participant Characteristics

Relative to the control group, the developmental dyslexia group had lower average scores for spelling abilities, reading comprehension, and reading speed (Table 1). However, the two groups were comparable in terms of age and intelligence (age: *t*(34) = 1.41, *p* = .167; IQ: *t*(34) = .50, *p* = .620).

### EEG Results

#### P120 and MMN Responses to Location Deviance

First, we examined whether developmental dyslexia affects ERP responses to location deviants (only standards and deviants presented on high-probability triplet endings were included in this analysis). Location deviants elicited a P120 followed by a location MMN, which was maximal at anterior frontal electrodes (Figure 2a-b). The location MMN appeared in both the dyslexia and control groups, but had a smaller amplitude in the dyslexia group. Compared with the standards, the location deviants elicited larger P120 responses in the control group, whereas the location deviants did not elicit larger P120 responses in the dyslexia group (see Appendix B for mean P120 and location MMN amplitudes in each condition). These observations were reflected in an ANOVA, indicating a significant interaction between sound location (standard, deviant) and group for the P120: *F*(1, 34) = 5.52, *p* = .025, η^2^p = .14; and the location MMN: *F*(1, 34) = 5.16, *p* = .03, *η*^2^p = .13; see the Appendix C for complete results. When analysing ERP responses separately in each group, location deviants elicited a significant location MMN in both the dyslexia group and the control group (*p* <.001; Figure 2c). Compared with the standards, location deviants elicited larger P120 responses in the control group (*p* = .011), but there was no significant difference in the dyslexia group (*p* = .49). Higher spelling scores on the Rechtschreibtest correlated with larger location deviance–induced P120 effects (*r* = .37, *p* = .03; see the Appendix C). No other correlations between language aptitude test scores and EEG responses were observed.

**Figure 2.**
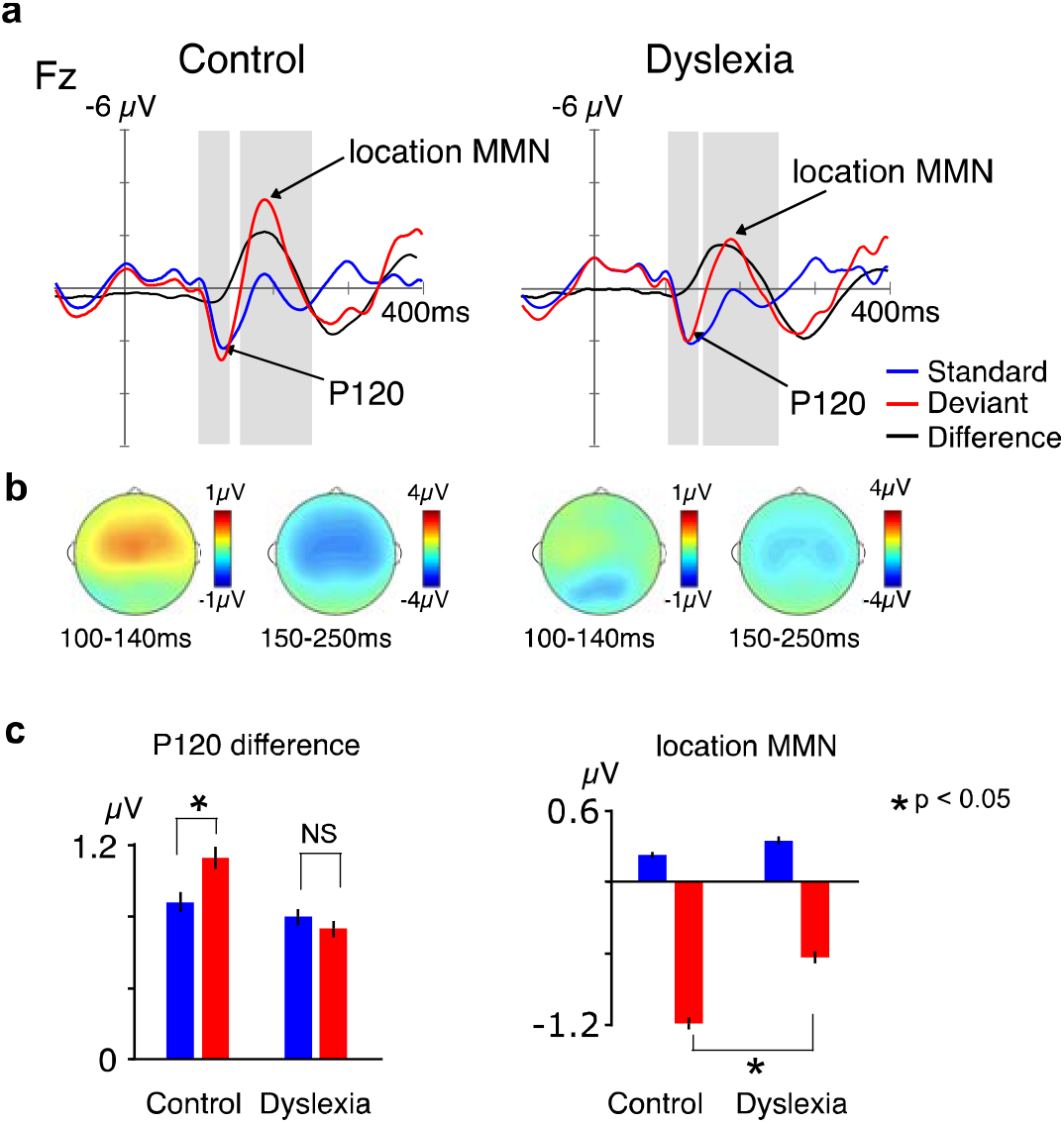
Location MMN results. **(a)** Mean ERP responses to triplet endings as recorded from the FZ electrode. Grey areas indicate the time windows used for quantifying the P120 (~100–140 ms) and location MMN (150–250 ms) components. Averaged ERP responses to location standard (blue) and deviant (red) triplet endings, as well as differences between them (black), are shown separately. **(b)** Isopotential maps showing the scalp distributions of differences between the ERPs evoked by location deviant triplet endings and those evoked by location standard triplet endings in the control (left) and dyslexia (right) groups. **(c)** Interactive effects of dyslexia and sound location at an anterior frontal region (average of F7, F5, F3, FT7, FC5, FC3, F1, FZ, F2, FC1, FCZ, FC2, F8, F6, F4, FT8, FC6, and FC4). Location deviant triplet endings elicited larger P120 amplitudes than location standard triplet endings in the control group, but this was not the case in the dyslexia group. Significant location MMN effects were observed in both groups but were larger in the control group than in the dyslexia group. Error bars indicate the standard deviation of the mean. Abbreviations: ERP, event-related potential; location MMN, location mismatch negativity.

#### P120 and MMN Responses to Statistical Deviance

Next, we examined whether developmental dyslexia affects ERP responses to statistical deviants (only triplet endings without location change were included in this analysis). Statistical deviants elicited a P120, followed by a sMMN, which was maximal over anterior frontal electrodes (Figure 3a-b; see the Appendix B for mean amplitudes of the P120 and sMMN components in each condition). Statistical deviants elicited a P120 in both groups, and an sMMN in the control group but not in the dyslexia group. These observations were reflected in an ANOVA, indicating a significant interaction between transitional probability (high vs. low probability), frontal vs. central, and Group: *F*(1, 34) = 5.50, *p* = .025, *η*^2^*p* = .14 (for complete results, see the Appendix D). Post-hoc tests revealed that the sMMN response at frontal electrodes was significant in the control group (*p* = .032) but not in the dyslexia group (*p* = .74; Figure 3c). At the central electrodes, the sMMN was not significant in both groups.

**Figure 3.**
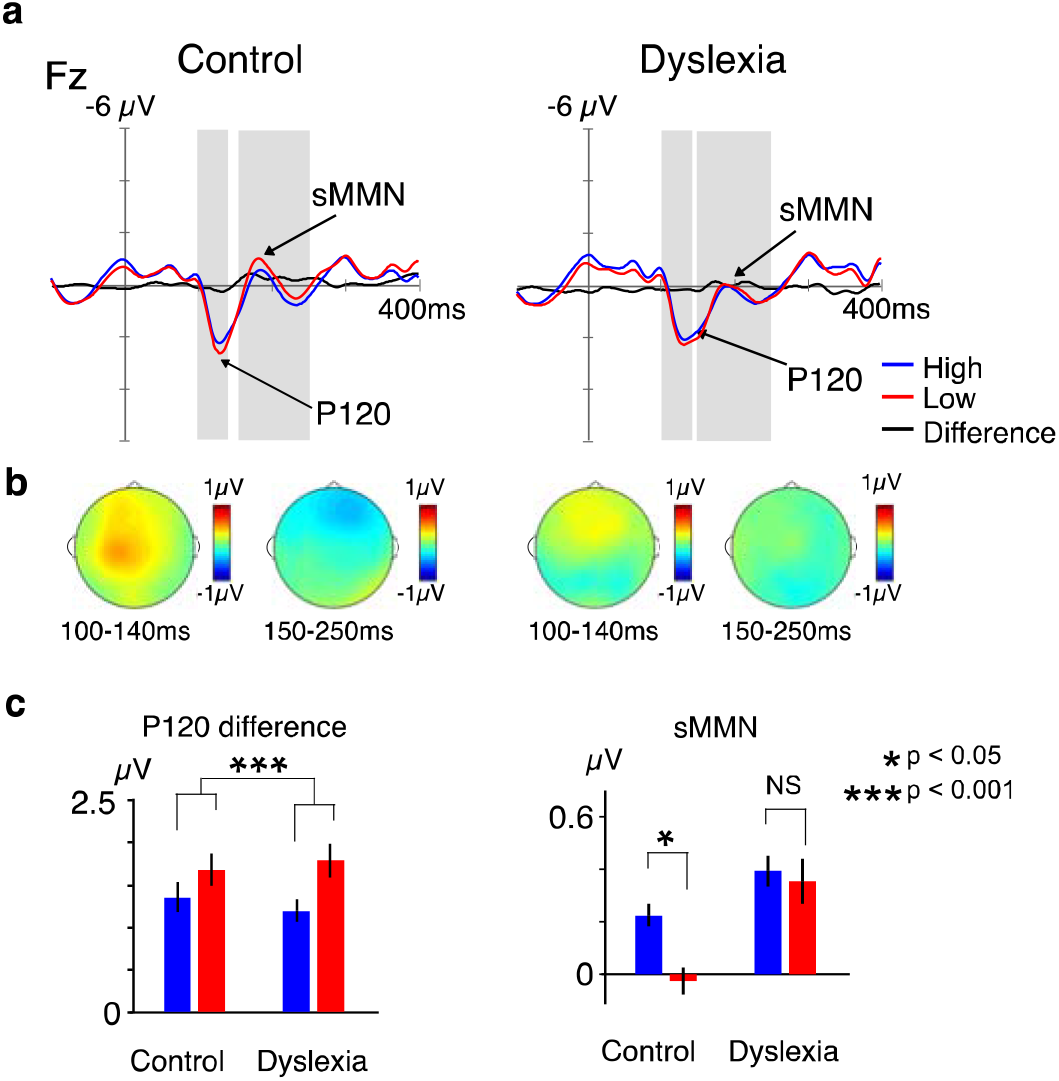
Statistical MMN results. **(a)** Mean ERP responses to triplet endings as recorded from the Fz electrode. The grey areas indicate the time windows used for quantifying the P120 (~ 100–140 ms) and sMMN (150–250 ms) component. Averaged ERP responses to statistically standard (blue) and deviant (red) triplet endings, as well as differences between them (black), are shown separately. **(b)** Isopotential maps showing the scalp distributions of differences between ERPs evoked by statistically deviant triplet endings and those evoked by statistically standard triplet endings. **(c)** Interactive effects of dyslexia and transitional probability at an anterior frontal region (average of F7, F5, F3, FT7, FC5, FC3, F1, FZ, F2, FC1, FCZ, FC2, F8, F6, F4, FT8, FC6, and FC4). In the medial electrodes, statistically deviant triplet endings elicited larger P120 amplitudes than the statistically standard triplet endings in both groups. Significant sMMN effects were observed in anterior frontal brain areas in the control group, but no sMMN effects were observed in the dyslexia group. Error bars indicate standard deviation of the mean. Abbreviations: ERP, event-related potential; sMMN, statistical mismatch negativity.

The effects of statistical learning on P120 amplitudes were prominent at medial electrodes (Figure 3b–c), as reflected by the significant interaction between transitional probability and lateralisation (F[2, 68] = 4.15, *p* = .020, *η*^2^*p* = .11). Post-hoc tests revealed that the P120 responses to statistical deviants were larger than those to the standards at medial electrodes (*p* = .009); however, no such effects were apparent in the left (*p* = .23) or right region (*p* = .53).

The effects of statistical learning on P120 amplitudes gradually increased as the experiment progressed to later blocks, as reflected by a significant interaction between transitional probability, lateralisation, and experimental blocks (*F*[4,136] = 2.78, *p* = .029, *η*^2^*p* = .08). Post-hoc tests revealed that the amplitudes of P120 responses to statistical deviants were larger in the third experimental block than in the first (*p* = .002) and second blocks (*p* = .004). Furthermore, in the analysis of ERP responses over the central region, the P120 response amplitudes elicited by statistical deviants were significantly different from those elicited by standards during the third block (*p* <.001); however, this was neither observed for the first (*p* = .15) nor the second block (*p* = .85).

### Behavioural Results

During the exposition phase, the participants discovered on average 94.3% *(SD =* .02) of the acoustical deviants (i.e., they showed a high performance in the cover task where participants had to detect high pitched tones). This indicates that the participants paid attention to the acoustical stimuli, while the task was relatively simple to carry out.

At the end of the exposition phase in each block, the participants listened to paired statistically deviant and standard triplets. A participant was asked to choose which triplet in the pair sounded more familiar and to rate their confidence in a given answer on a scale ranging from 1 (no certainty) to 5 (certainty). To evaluate the performance in the behavioural testing phase, we used a two-tailed t-test to determine whether the frequency of correct answers during behavioural testing exceeded chance levels (i.e., *p >* .05). Further, we used ANOVAs to compare response accuracies, RT, and confidence ratings between dyslexics and controls. ANOVAs were computed with Group (dyslexia vs. control) as a between-subjects factor and experiment block as a within-subject factor (three blocks; the first, second and third blocks, instead of the actual six blocks of the experiment to obtain a higher signal-to-noise ratio).

The two-tailed t-test revealed the frequency of correct answers was significantly higher than chance levels in the control group (*p* = .017, *Cohen’s* d = .325) but not the dyslexic group (*p* = .520). ANOVAs revealed no effects on response accuracies and confidential rating in both the dyslexia and control groups (Figure 4, and see Appendix E). As for the reaction time, a significant main effect of the experimental block was noted (*F*[2, 68] = 7.92, *p* <.001, *η*^2^*p* = .189). Post-hoc tests revealed that the reaction time in the second and last blocks was significantly faster than that in the first block (2nd: *p* = .006; 3rd: *p* <.001). No other effect was found in the ANOVA of behavioural results.

**Figure 4.**
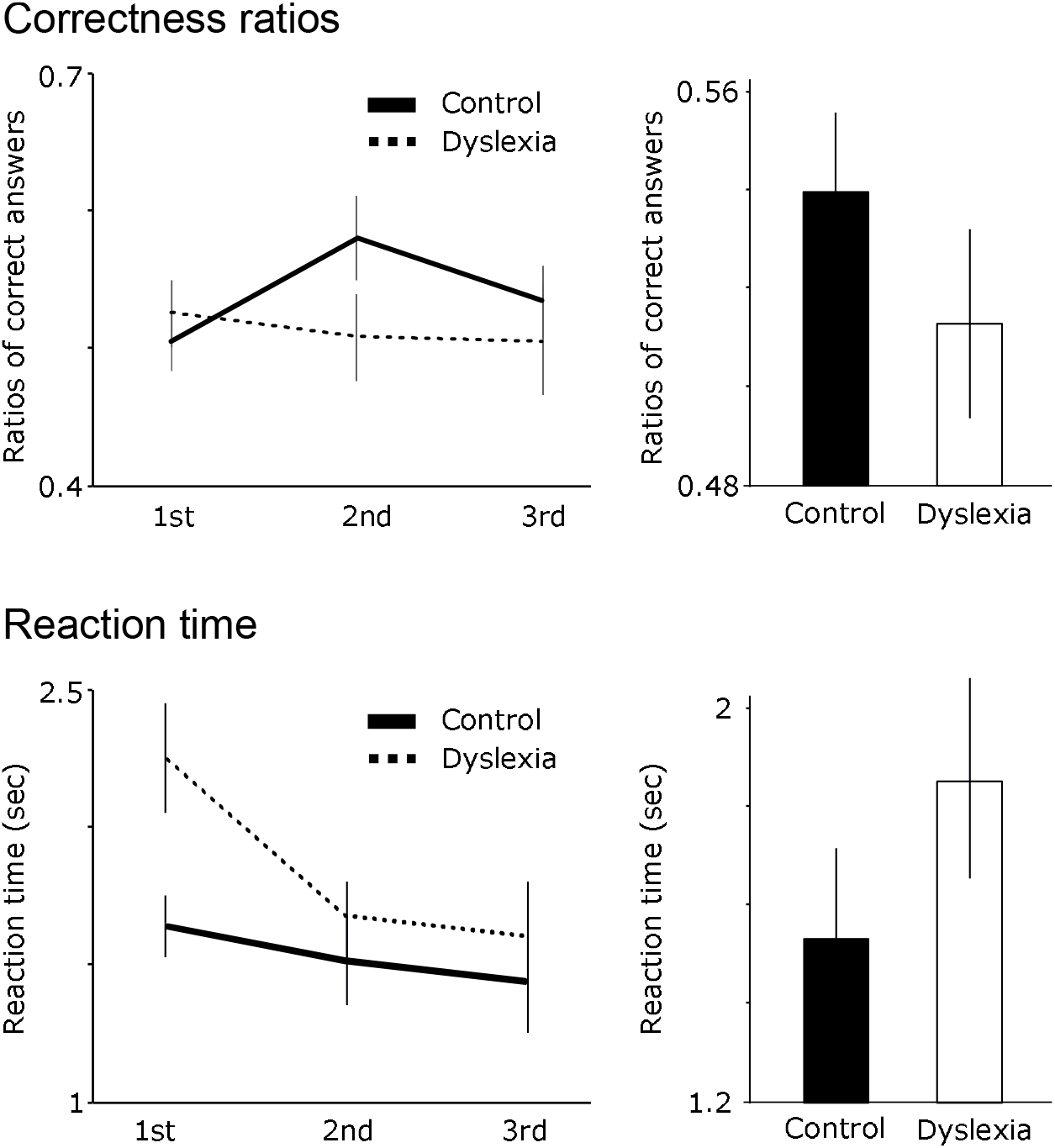
Behavioural testing results across experimental blocks. Error bars indicate standard deviations. The ANOVA revealed no significance difference in correctness ratios and reaction time between dyslexia and control groups.

To determine whether response accuracy rates correlated with confidence ratings, we also conducted a Pearson correlation analysis. Response accuracies did not correlate with confidence ratings in both the control (*r* = −0.18, p = .19) and dyslexia (*r* = −0.05, *p* = .70) groups. Similarly, response accuracies did not correlate with reading and spelling test scores.

## Discussion

In this study, we assessed the relationship between developmental dyslexia and ERP responses to a continuous stream of sound triplets in which some triplet endings were either location deviants or statistical deviants. We found that statistical deviants elicited small sMMN and that location deviants elicited a large location MMN. Compared with controls, the participants with developmental dyslexia exhibited a smaller location MMN and no sMMN. Whereas there was a tendency to a higher proportion of of correct responses and decreased reaction times in the control compared to the dyslexia group,there was no significant between-group difference for those two variables, reflecting the ability to learn the statistical characteristics of stimuli. Our findings indicate that the neural functions underlying the sMMN and the location MMN are impaired in individuals with developmental dyslexia. This is in agreement with previous evidence of a smaller MMN in developmental dyslexics (Gu & Bi, 2020; Lachmann et al., 2005; Neuhoff et al., 2012). Our findings also suggest that at the group level, the MMN may be more sensitive to mild difficulties than behavioural measures.

The location MMN is a response reflecting auditory perceptual memory operations that are instantly updated after new information is obtained (Bendixen, Prinz, Horváth, Trujillo-Barreto, & Schröger, 2008; Sussman & Winkler, 2001). The sMMN, on the other hand, relies on memory representations that are formed from the implicit knowledge of sequential statistical structure (i.e., knowledge of stimulus transitional probabilities; cf., e.g., Koelsch et al., 2016; Tsogli et al., 2019). Notably, the sMMN effects elicited in our experiment were maximal at anterior frontal electrodes (Figure 3b); however, the location MMN effect was more broadly distributed from central to frontal areas (Figure 2b). This suggests the possibility that neural sources of the sMMN are different from those of the location MMN. However, our findings showed that people with dyslexia also exhibited smaller location MMN.

A past study found a diminished pitch MMN but an enhanced location MMN in dyslexia (Kujala et al., 2006). We assumed that the combination of location and statistical MMN in our paradigm may lead to different results from Kujala’s study.

Another possibility is that the acoustic perceptual dysfunction of dyslexic individuals also influenced neural processing of a sMMN. For example, past studies have showed faster decay of dyslexics’ perceptual memory trace behaviorally (Jaffe-Dax et al., 2017; Lieder et al., 2019) and in brain activity (Jaffe-Dax 2015, 2017), specifically in the auditory cortex (Gertsovski & Ahissar, 2022 Perrachione et al., 2016). They proposed that it reduces learning of complex rules (e.g. Virtala et al., 2021) and particularly hampers acquisition of categories (Gertsovaki & Ahissar, 2022).

Both location MMN and sMMN were preceded by P120 responses. Compared with the controls, the participants with developmental dyslexia exhibited no difference in P120 responses between location deviants and standards. Previous studies have reported the atypicalities of the P1 component in sound processing and dyslexia (Stefanics et al., 2011) and implicit and statistical learning (Jaffe-Dax et al., 2017). The P120 found in our study may be a P1-like component in terms of the positive component temporally adjacent to the N1 or MMN effects. Further, given the fundamental differences in the memory system between ERPs elicited by statistically deviant stimuli and location deviants as stated above, the P120 components elicited by statistically deviant stimuli may also be distinguishable from those of the P120 components elicited by location deviants. However, considering the difference in the time window and that this effect was not hypothesised, it will not be discussed further until future studies replicate this effect. Further studies are therefore necessary to elucidate the neural basis underlying P120 components.

It has recently been considered that a domain-general statistical learning impairment, rather than a specific impairment in phonological processing (Ramus et al., 2003; Vellutino et al., 2004), may underlie developmental dyslexia. For example, individuals with developmental dyslexia show weaker domain-general statistical learning across sensory domains (Hung, Frost, & Pugh, 2018), such as auditory (Arciuli & Conway, 2018; Dobó et al., 2021; Gabay et al., 2015; Kahta & Schiff, 2019;

Vandermosten et al., 2019) and visual stimuli (Sigurdardottir et al., 2017). Furthermore, statistical learning impairment in developmental dyslexia is not limited to speech stimuli but also occurs in non-speech stimuli (Plakas, van Zuijen, van Leeuwen, Thomson, & van der Leij, 2013). Our results support these past findings; our study explored effects of deviation from predictions based on physical and statistical properties using non-speech auditory stimuli, and our findings showed that statistical learning processes and basic auditory processes are affected in individuals with developmental dyslexia.

Some new theoretical frameworks have been proposed to explain statistical learning impairment in dyslexia. For example, a multicomponent memory network, referred to as the Statistical Learning and Reading (SLR) model (Lee et al., 2022), represents a domain-general system that consists of a short-term memory subsystem, an explicit declarative long-term memory subsystem, and an implicit procedural long-term memory subsystem. The two long-term memory subsystems are distinguishable in terms of the attentional demands required for encoding and storage of information, with more controlled attention (i.e., a top-down selective attention to learning stimuli) than automatic attention (i.e., a bottom-up involuntary attention to salient stimuli) needed in the explicit declarative subsystem, and, conversely, more automatic attention than controlled attention needed in the implicit procedural subsystem. The model output is a statistically optimal representation as manifested by the neural and behavioural response of statistical learning and reading activities.

In this study, participants conducted the behavioural testing at the end of each block. In this experimental paradigm, it would be possible that they tended to pay more attention to the tone sequence in the later blocks since they anticipated they may experience a test later on. However, it is also important to see the time course of statistical learning effects. Indeed, this study detected that participants did not show statistical learning effects in the first block and a difference between groups. However, the difference gradually became larger particularly in the second block whereafter it stayed at about the same level (Figure 4, top). Further, the control group but not the dyslexic group showed an above-chance level in behavioral performance of statistical learning. However, no significant group difference may suggest the possibility that the two groups learned the sequence equally. Further research is necessary to reveal why and how the MMN is more sensitive to group difference than behavioural measures.

In conclusion, our findings lay forth evidence that location deviants elicit a distinct classical MMN, which was larger in controls than in individuals with developmental dyslexia. Statistical deviants elicited a sMMN in controls, whereas in individuals with developmental dyslexia, sMMN was not recognizable. Our findings add to and are consistent with the so far scarce evidence showing that statistical learning and the underlying neural correlates may be impaired in individuals with developmental dyslexia. Thus, exploring those neural correlates may contribute to a better understanding of the cognitive processes underlying the acquisition of the rules and regularities that guide the arrangement of elements in ordered sequences such as language and music.

## Acknowledgements

This study was supported by the German-Israeli Foundation for Scientific Research and Development and World Premier International Research Centre Initiative (WPI), MEXT, and JSPS KAKENHI Grant Number 20K22676, Japan. The sponsors played no role in the study design; the collection, analysis, and interpretation of the data; or the writing of the manuscript

## Extended data

**Appendix B. Mean ERP amplitudes under different conditions.** ERP, event-related potential.

**Appendix C. ANOVA results for the effects of location deviance.** (a) Correlation analysis and ANOVA results for P120 component amplitudes and (b) ANOVA results for location MMN effects. Location MMN, location mismatch negativity.

**Appendix D. ANOVA results for the effects of statistical deviance.** (a) ANOVA results for P120 component amplitudes and (b) sMMN effects. sMMN, statistical mismatch negativity.

**Appendix E. ANOVA results for data from a behavioural test assessing the statistical learning of stimulus characteristics.**

